# RNA-seq transcript quantification from reduced-representation data in recount2

**DOI:** 10.1101/247346

**Authors:** Jack M. Fu, Kai Kammers, Abhinav Nellore, Leonardo Collado-Torres, Jeffrey T. Leek, Margaret A. Taub

## Abstract

More than 70,000 short-read RNA-sequencing samples are publicly available through the recount2 project, a curated database of summary coverage data. However, no current methods can be directly applied to the reduced-representation information stored in this database to estimate transcript-level abundances. Here we present a linear model taking as input summary coverage of junctions and subdivided exons to output estimated abundances and associated uncertainty. We evaluate the performance of our model on simulated and real data, and provide a procedure to construct confidence intervals for estimates.

## Background

RNA sequencing (RNA-seq) can be used to measure gene (and transcript) expression levels genome-wide. Large-scale RNA-seq datasets have been produced by studies such as the GTEx (Genotype-Tissue Expression) consortium [1], which comprises 9,662 samples from 551 individuals and 54 body sites (under version 6), and the Cancer Genome Atlas (TCGA) study [2], which comprises 11,350 samples from 10,340 individuals and 33 cancer types. Furthermore, public data repositories such as the Sequence Read Archive (SRA) host tens of thousands of human RNA-seq samples [3]. These data collectively provide a rich resource which researchers can use for discovery, validation, replication, or methods development.

These data are even more valuable when processed in a consistent manner and presented in an accessible format. Researchers can query the database to test any relevant hypotheses they may have. The recently published recount2 project [4] is the result of such an undertaking. All raw data from the thousands of sequencing studies were aligned to a common reference genome using a scalable and reproducible aligner Rail-RNA [5]. Summary measures (gene, exon, junction, and base-pair level coverage) were derived from the Rail-RNA output and made available in an R package and through a web portal (https://jhubiostatistics.shinyapps.io/recount/). The over 70,000 curated samples have reads whose lengths fall within approximately 5 distinct peaks and were either paired or unpaired (see **Table 1**)

**Table 1:** The number of samples in recount2 that fall closest to each read length by paired-status category. 5 distinct peaks of read lengths (37, 50, 75, 100, and 150bp) were observed in recount2, and samples are assigned to the closest matching read length out of the 5 above categories.

| Read length | 37 | 50 | 75 | 100 | 150 |  |
| --- | --- | --- | --- | --- | --- | --- |
| Single end | 5,714 | 10,557 | 2,395 | 3,459 | 232 | 22,357 |
| Paired end | 1,953 | 14,123 | 14,965 | 16,725 | 874 | 48,640 |
|  | 7,667 | 24,680 | 17,360 | 20,184 | 1,106 | 70,997 |

Currently, recount2 provides summary measures that directly allow for analyses like annotation-agnostic base-pair level and annotation-specific gene/exon/junction differential expression. However, transcript-level abundance estimates are missing from recount2, preventing subsequent transcript-level analyses. Despite the existence of many successful transcript quantification programs (such as Cufflinks [6], Kallisto [7], Salmon [8], and RSEM [9]), this deficiency persists because methods capable of estimating transcript abundances using the summarized output collected in recount2 do not exist. Here, we present a linear model-based method to accomplish this estimation task.

Previous linear model-based transcript abundance estimation techniques include IsoformEx [10], MultiSplice [11], and CIDANE [12]. IsoformEx transforms aligned reads into RPKM (reads per kilobase of transcript, per million mapped reads) of splice junctions and disjoined exons to apply a length-weighted non-negative least squares regression for abundance estimation. In addition to the basic exon and junction counts, MultiSplice and CIDANE account for reads that are more identifiable to a unique transcript such as reads that span multiple junctions, or a read-pair that uniquely links multiple exons. Since our model does not have access to the highly discriminating read-level information leveraged by MultiSplice and CIDANE, we were restricted to maximizing the summary coverage statistics accessed by IsoformEx. Toward that goal, our model further subdivides exonic segments and introduces an aligner-estimated model matrix. Our objective in developing this method was not to be faster or more accurate than existing methods operating on raw sequencing data, but to provide more full utilization of the data in the recount2 project through transcript-level abundance estimates.

## Results

### Overview of method

recount2 includes a repository of coverage summary measures, including coverage of exon-exon splice junctions, produced by a uniform application of the aligner Rail-RNA to more than 70,000 publicly available RNA-seq samples. For a given read length and a reference transcriptome, we determine a set of sufficient **features** comprised of subdivided exonic segments and exon-exon junctions, such that the coverage of these features adequately summarizes the transcript quantification encoded in the raw reads. Counts of reads overlapping the features are the sufficient statistics of our linear model, which we denote as **feature counts** (see **Figure 1**).

**Figure 1:**
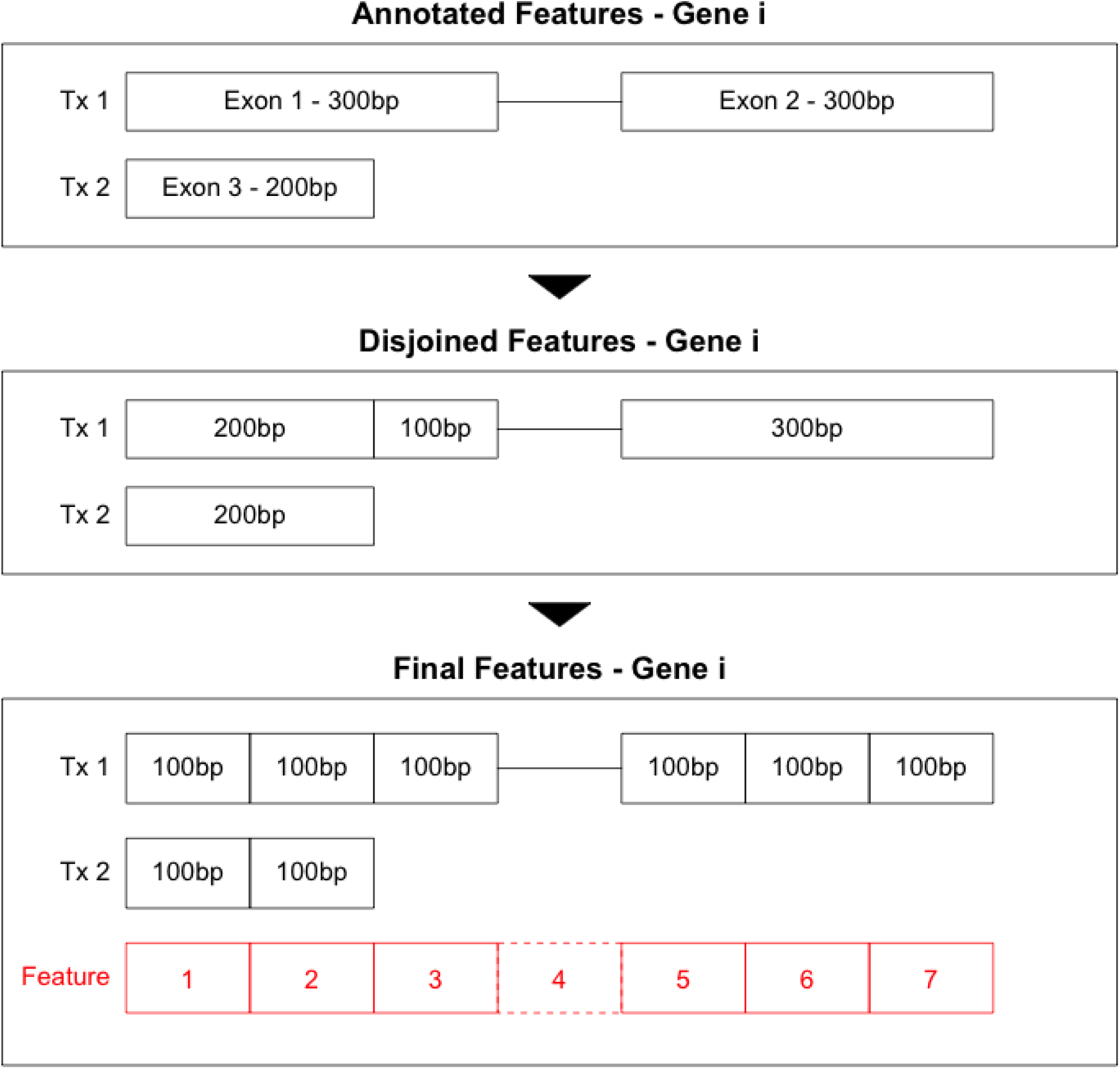
Given the read length of a particular experiment and a reference transcriptome, we determine a set of sufficient **features** such that the coverage of these features adequately summarizes the transcript quantification encoded in the raw reads. This figure illustrates the process to determine the set of features for an example gene with 2 transcripts and 100bp reads. We first disjoin the annotated exons into non-overlapping bins. Any remaining exonic segments longer than twice the read length are further evenly subdivided to be below 100bp. Each unique splice junction is included without modification as a feature. The number of reads overlapping the final set of features are the sufficient statistics for our linear model and serve as a compression of the 18 raw read-level data.

Using these feature counts as the dependent variable, we fit a non-negative least squares regression model to estimate the underlying transcript abundances. The independent variables in our model are transcriptome annotation-specific, and are denoted as **feature probabilities**. A feature probability encodes the chance that a random read from a transcript will contribute an observed count to the corresponding feature. Standard error estimates are reported that reflect our model’s confidence in abundance assignment. Lastly, our method also reports a uniqueness score for each transcript that reflects how distinguishable each transcript is compared to other transcripts during quantification. Further details about our methods are described in Methods and are implemented in the R package recountNNLS available from https://github.com/JMF47/recountNNLS.

### Performance on Dirichlet-negative binomial simulated data

Using simulated data based on a Dirichlet-negative binomial specification described in Methods, we evaluated the performance of our model and the commonly-used pipelines HISAT2-Cufflinks [13, 6], Kallisto [7], Salmon [8], and RSEM [9]. We simulated reads from 18,303 transcripts using all annotated features on chromosomes 1 and 14, including 10 scenarios of varying readlength and paired-end status, using the polyester R package [14]. Our method was run on the reduced-representation output from applying the aligner Rail-RNA [5] to the simulated FASTA files. All other methods extracted information from the full simulated FASTA files. The accuracy of all methods across the range of simulated scenarios is measured in mean absolute error of estimated abundance compared to the truth, and is visualized in **Figure 2** with numbers reported in **Supplementary Table 1**.

**Figure 2:**
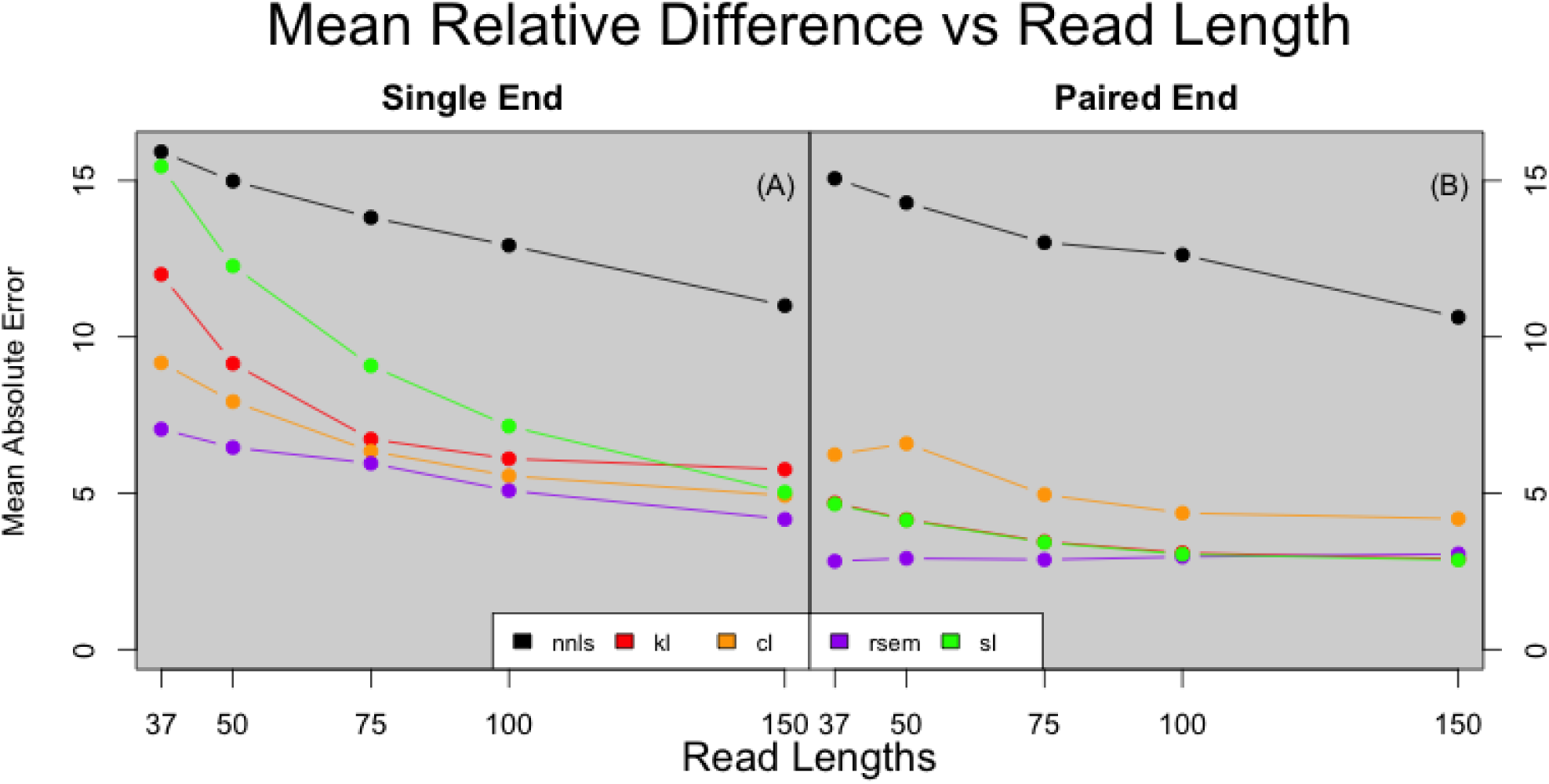
The mean absolute error (MAE) of each method’s performance plotted over read length in Dirichlet-negative binomial simulations, separated into panels by paired-end status. Colors correspond to method. In both single and paired-end simulations, all methods were able to improve performance as read length increases. Kallisto (kl), HISAT2-Cufflinks (cl), Salmon (sl), and RSEM (rsem) are able to leverage information encoded by the pairs of reads to further reduce error, whereas our method (nnls) is not. The effective read length under paired-end reads is a combination of the insert size and sequencing read length.

**Figure 3** shows results from the 75bp, paired-end simulation scenario described in Methods and helps illustrate the utility of the uniqueness score produced by our model. The distribution of uniqueness scores is visualized in **Figure 3 (A)**. **Figure 3 (B)** indicates that bias in transcript estimates decreases as uniqueness scores increases. In **Figure 3 (C)**, we observe that as uniqueness scores decrease, the standard errors reported by our model increase to reflect the uncertainty caused by similarity between transcripts.

**Figure 3:**
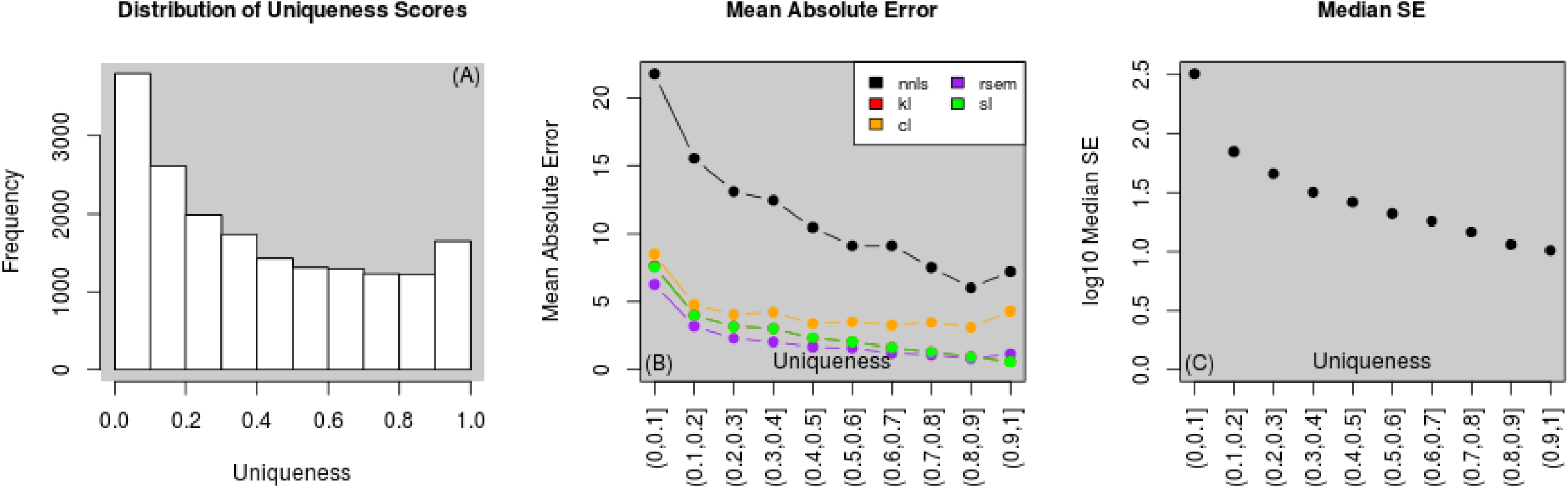
For the 75bp, paired-end Dirichlet-negative binomial simulation, our model produces uniqueness scores to reflect our ability to distinguish between transcripts during modeling **(A)**. Panel **(B)** demonstrates that as the uniqueness of a transcript increases, our method is able to improve in bias as measured by mean absolute error. Colors correspond to method which are denoted as NNLS (nnls), Kallisto (kl), HISAT2-Cufflinks (cl), Salmon (sl), and RSEM (rsem). Panel **(C)** depicts a decrease in median standard errors reported by our model as uniqueness increases.

### Performance of confidence intervals

To assess confidence interval coverage probabilities, for a random subset of 2000 transcripts from chromosome 1, we simulated 100 datasets for each of the 10 Dirichlet-negative binomial scenarios. For each dataset, we constructed 95% confidence intervals and evaluated whether those intervals overlapped the truth. We observe that as uniqueness score of a transcript increases, confidence intervals constructed for that transcript are more likely to capture the truth at least 95 times out of the 100 repeats. For example, for 75bp single-end reads, around 85% of transcripts in the lowest uniqueness category achieve nominal significance, compared to around 96.5% in the highest uniqueness category. We also observe that as read length increases, the proportion of transcripts with 95% confidence intervals that overlap the truth in at least 95 out of the 100 repeats generally increases. (**Figure 4 (B, C)**).

**Figure 4:**
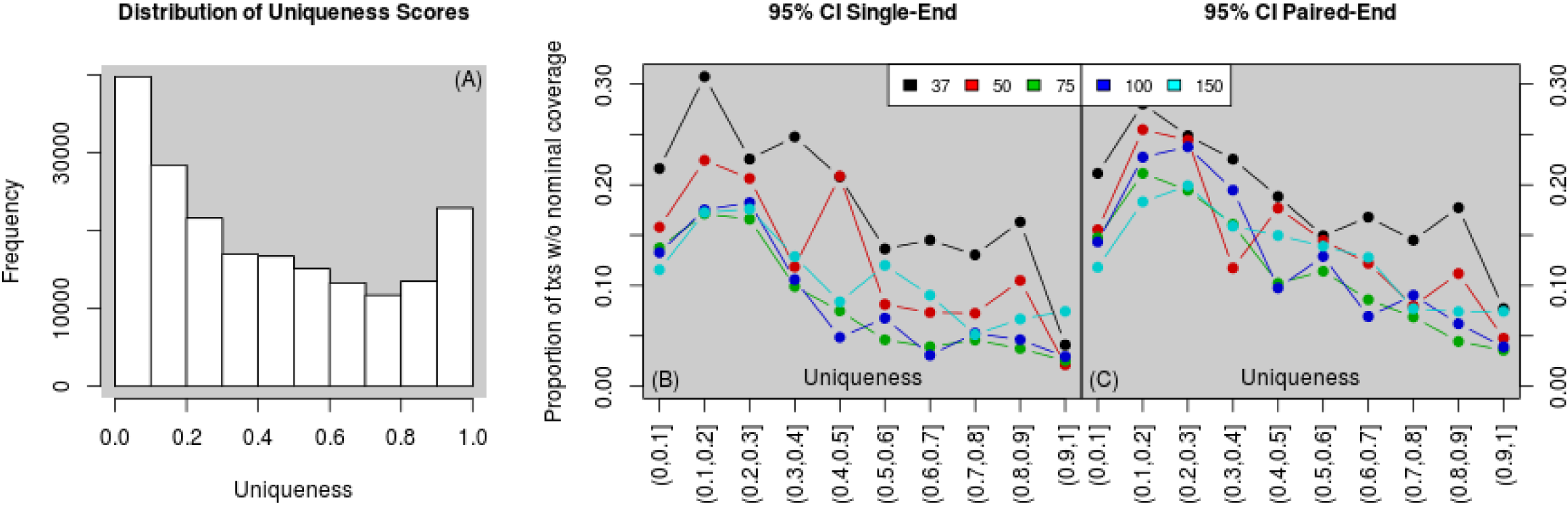
For 2000 randomly chosen transcripts on chr1, we simulated 20x coverage 100 times for each of the 10 Dirichlet-negative binomial scenarios. Panel **(A)** presents the distribution of uniqueness scores for each of the 2000 randomly chosen transcripts. For each transcript, we constructed 95% confidence intervals across the 100 repeats, and made note if the set of intervals overlapped the truth in less than 95 of the 100 repeats. Panel **(B)** visualizes how often a transcript in a given uniqueness score range was noted in the step previous as having intervals overlapping the truth in less than 95 of 100 repeats for single-end scenarios, while **(C)** visualizes the same information for the paired-end scenarios. Colors correspond to read lengths.

### Performance on hybrid simulated data

To assess performance on simulated data that might more accurately reflect biology than the Dirichlet-negative binomial scenario described above, we also simulated a 75bp, paired-end dataset using the RSEM estimated transcript abundances for the ERR188410 sample of the GEUVADIS Consortium data as ground truth expression levels. RSEM was excluded from this comparison. Our method’s performance relative to the others is consistent with our above Dirichlet-negative binomial simulation of 75bp, paired-end reads (**Supplementary Table 1, last row**). The distribution of uniqueness scores is seen in **Figure 5 (A)**. Our method again shows decreasing mean absolute error accompanying a rise in uniqueness score **Figure 5 (B)**. The median standard error decreases with increasing uniqueness scores in **Figure 5 (C)**.

**Figure 5:**
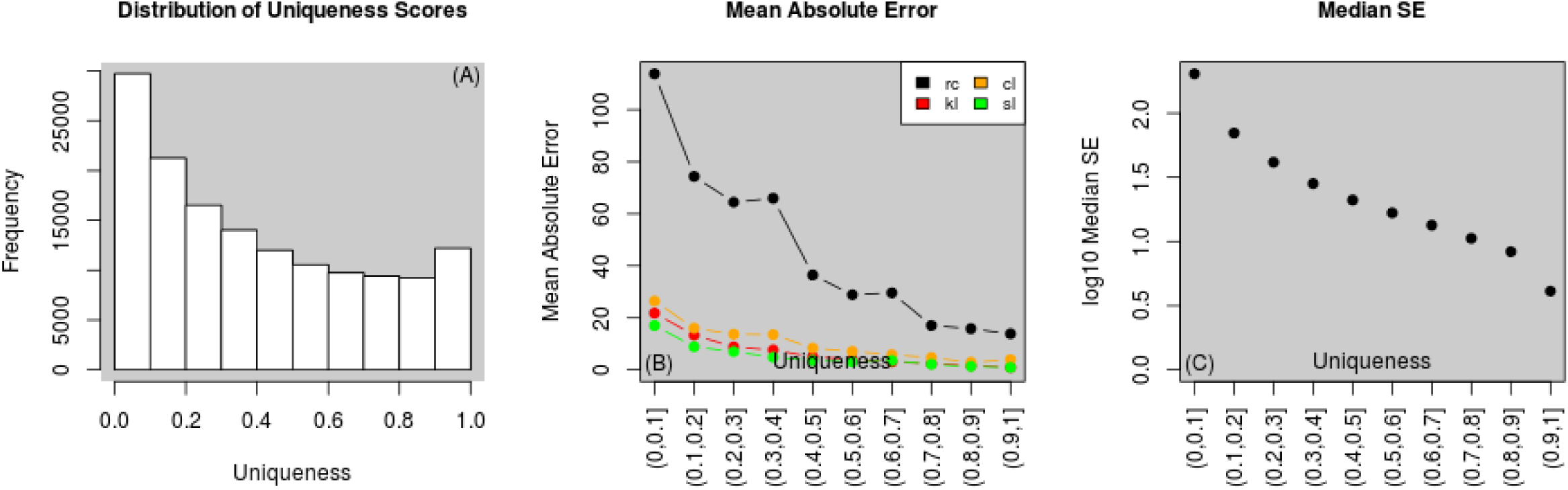
This figure demonstrates the output for the hybrid simulation scenario, where ground truth is the estimated counts from applying RSEM to ERR188410 of the GEUVADIS Consortium dataset, a 75bp paired-end experiment. **(A)** Displays the distribution of uniqueness scores used in our linear estimation. Panel **(B)** demonstrates that as the uniqueness of a transcript increases, our method is able to improve in bias as measured by mean absolute error. Panel **(C)** depicts a decrease in median standard errors reported by our model as uniqueness increases.

### Performance on GEUVADIS Consortium data

Using recount2 feature counts of sample ERR188410 (a 75bp paired-end sample) from the GEUVADIS dataset [15], we ran recountNNLS to estimate transcript abundance levels. We also downloaded the FASTQ files for this sample and applied the 4 other methods mentioned above to estimate transcript abundances. Pair-wise comparisons of the estimates were carried out to evaluate Spearman’s correlation and concordance of transcripts assigned non-zero expression.

For this sample, the Spearman correlations between the other methods are high, at greater than recountNNLS achieves much more moderate correlation of approximately 0.65 with the other methods. The lower triangle of **Table 2** presents the number of transcripts that the corresponding methods both assigned non-zero expression to. Salmon [8] detected the most transcripts with nonzero expression at 83,733, while our method reported the least with 68,568 transcripts.

**Table 2:**
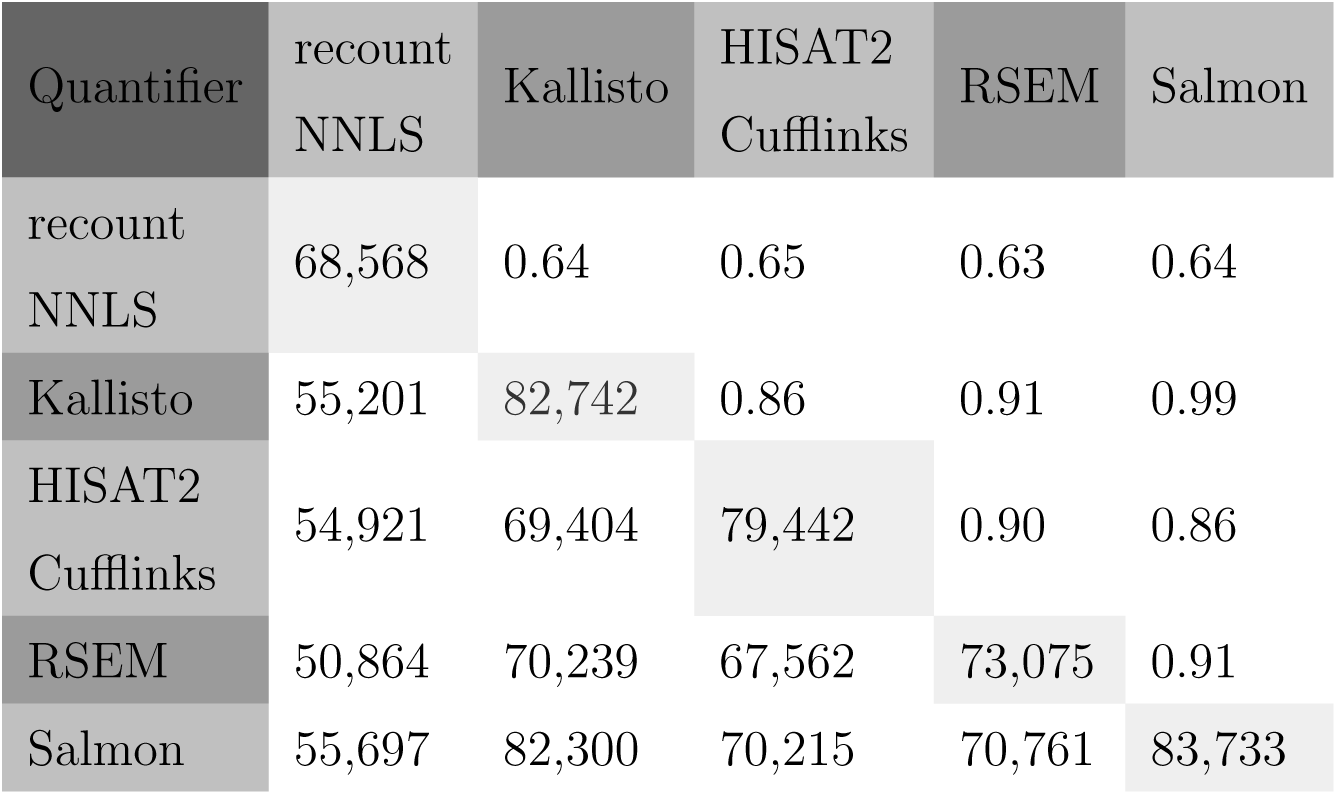
Pair-wise comparison of the evaluated methods on example ERR188410 of the GEUVADIS consortium dataset. The upper half shows pair-wise Spearman’s correlation. The lower half shows the number of transcripts that both methods assigned non-zero expression to. The diagonal shows the number of transcripts that each individual method assigned non-zero expression to.

## Discussion

We have presented here a method to provide transcript-level abundance estimates on the reduced-representation expression data available in recount2. Our model’s performance most closely approximates other methods in the 37bp single end read setting. All methods are able to improve as read lengths increase (**Figure 2**), likely because longer reads have a higher probability of being uniquely attributable to a single transcript. However, our method is not able to leverage the full information of longer reads, such as reads that span a unique sequence of junctions, and showed more modest improvements. Similarly, for paired-end scenarios, the insert length works in conjunction with the read length to dramatically increase the probability of uniquely assigning a read for those methods who have access to such information. Still, given the unique resource that recount2 represents, including transcript-level abundances will greatly increase the potential downstream uses of the data in this repository.

In order to examine loci where our model achieves higher or lower accuracy, we develped a measure of transcript uniqueness. Many loci have annotated transcripts that are structurally very similar. Unsurprisingly, expression levels for transcripts that are highly similar are difficult to tease apart. To identify how structurally similar one transcript is among a set of transcripts, in the context of linear estimation, we calculate a uniqueness score that represents the proportion of variability in feature probabilities of a transcript that can be explained by the other transcripts. This score ranges from 0 to 1, with 0 indicating that a transcript’s feature probabilities can be perfectly recapitulated by other transcripts and 1 indicating that a transcript is wholly unique. As noted above, we see a clear relationship between the uniqueness score and the estimated accuracy of our method in both **Figure 3 (B)** and **Figure 5 (B)**. The other methods evaluated also tend to show a trend in decreasing bias with increasing uniqueness score from our model.

Our model’s standard error estimates are also related to the uniqueness score of a transcript. Under our Dirichlet-negative binomial and hybrid simulations, we observe an increase in median standard errors as uniqueness decreases in **Figure 3 (C)** and **Figure 5 (C)**. Similarly, in **Figure 6**, we observe the structure and estimates of 2 selected genes from sample ERR188410 of the GEUVADIS Consortium dataset. For the gene KLHL17, all 5 transcripts have unique features that make the transcripts highly distinct. Our method shows that the standard errors are relatively low (**Figure 6, top**). In the gene G6PD, there are strong structural similarities between some of the transcripts. The difference in the identifiability of the transcripts is clearly reflected in our reported standard errors: transcripts that are difficult to distinguish from others are assigned higher standard errors (**Figure 6, bottom**). In particular, the top two transcripts are almost identical, with uniqueness scores of far less than 0.01 and inferred standard errors orders of magnitude larger than other transcripts at this locus.

**Figure 6:**
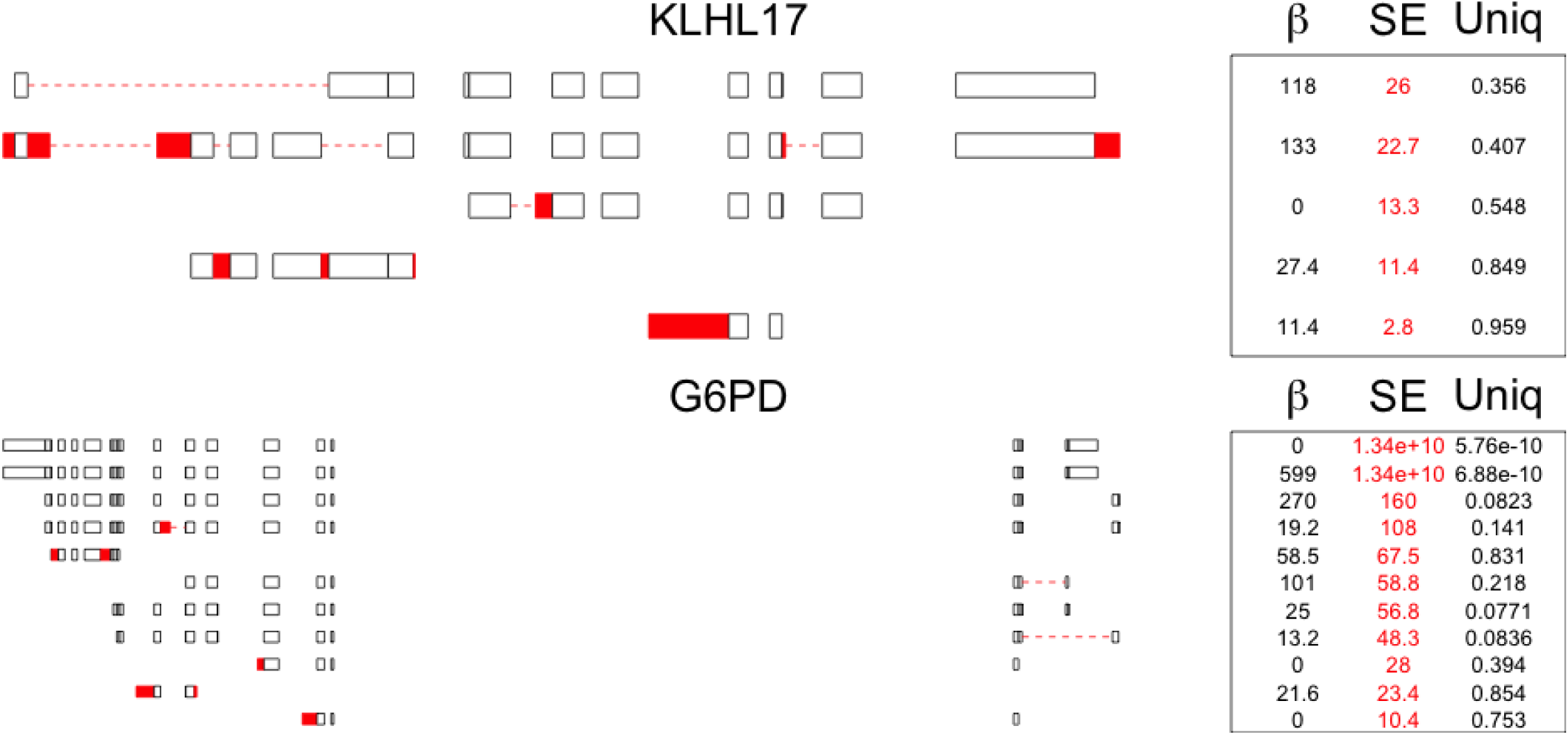
The gene on top (KLHL17, ENSG00000187961.13) is comprised of 5 transcripts, while the gene on the bottom (G6PD, ENSG00000160211.15) consists of 8 transcripts. For each panel, the transcript structure is shown on the left, with unique features highlighted in red. On the right side of each subplot are the estimated abundances in black and the standard errors of these estimates in red, on the scale of number of fragments. The transcripts are ordered by decreasing standard error from top to bottom. Uniqueness scores for each transcript are also reported. The higher estimated standard errors for the top transcripts in G6PD reflect the sequence similarity between those transcripts.

The more uniquely identifiable a transcript is, the more likely that transcript will have confidence intervals that cover the truth in greater than 95 out of 100 repeated simulations (**Figure 4 (B,C)**). Transcripts of low uniqueness tend to have higher bias and less-than-nominal confidence interval coverage. Similarly, as read length increases, uniqueness scores of all transcripts tend to improve overall. Thus the validity of our confidence intervals also improves as read length increases.

We have already run our model on all data currently in the recount2 database and have uploaded the results for public access. Working with the transcript abundances produced by our method is straightforward. For a given SRA project id (x) currently curated in recount2, one can access the transcript quantification stored as a RSE object by installing the recountNNLS R package and calling a single function, getRseTx(x). We also include an example differential transcript expression analysis of healthy versus cancer TCGA breast samples in the **Supplementary Materials**. We input our model estimates into a popular differential expression pipeline using the R packages limma [16] and edgeR [17] to produce estimates of transcript-level differential expression between these groups of samples.

Our model is readily able to adjust for factors that might affect quantification (such as GC content, mappability, and transcript location bias) by adjusting the feature probability matrices. For example, to adjust for GC content, we could learn the GC content bias of the sample by selecting for the subset of 1-transcript genes and assessing GC bias using their sequence composition and expression levels. The selected transcripts can be broken down into the set of features and feature counts that they are comprised of. Using a loess smoother, one could model the relationship between the GC content of those features and the feature counts. This relationship could then be used on multi-transcript genes to up-weight or down-weight the feature probability matrix entries. Substituting the adjusted matrices into NNLS estimation would yield GC-adjusted estimates. Similar processes can be carried out for any kind of adjustment for which one could attain feature-level characteristics, such as mappability, positional biases, etc.

## Conclusion

We have presented a method for estimating transcript-level abundances and standard errors, as well as a procedure to construct confidence intervals, based on a set of reduced representation data on more than 70,000 RNA-seq samples in recount2. The quantified transcript abundances for the more than 70,000 samples of recount2 are now available for direct access and download via both the R package recount and the recount2 website https://jhubiostatistics.shinyapps.io/recount/.

## Methods

### recount2 summary measures

recount2 includes a repository of coverage summary measures produced by a uniform application of the aligner Rail-RNA to more than 70,000 publicly available RNA-seq samples. For each sample, recount2 contains two primary files necessary for our linear modeling approach. First, each sample has a BigWig-format file [18] containing the number of reads that overlap each genomic position of the hg38 assembly. Secondly, each sample has a file containing the number of reads spanning observed exon-exon splice junctions. Other useful summarizations are also available directly from recount2, like pre-computed exon-level and gene-level coverages based upon the GencodeV25 reference transcriptome using the above mentioned BigWig files.

### Sufficient statistics for transcript quantification

Given the read length of a particular experiment and a reference transcriptome, we determine a set of sufficient **features** such that the coverage of these features adequately summarizes the transcript quantification encoded in the raw reads. For simplicity, we illustrate our definition of features with an example gene containing two transcripts, and with an example data-generating experiment with read lengths of 100 base pairs, but our method generalizes to arbitrary transcript structure and different read lengths.

Consider the gene portrayed in **Figure 1**, which is composed of 2 transcripts, 3 distinct exons, and 1 exon-exon junction, and suppose that the experiment produces reads of length 100bp. We first disjoin the annotation into unique, non-overlapping sub-exonic segments, similar to the scheme that IsoformEx [10] employs. However, in a process unique to our model, any bins longer than 200bp (twice the experiment read length) are then further evenly subdivided so that the largest resulting piece is less than 100bp. This process increases the identifiability of the transcripts. For our example, the final product is a set of 7 features, of which 6 are sub-exonic segments while 1 is an exon-exon splice junction.

The sufficient statistics for our linear model are the counts of reads that overlap each feature, which we will denote as **feature counts**. To extract the feature counts given a set of features, we query the BigWig files for the coverage of sub-exonic sections, and the junction file for the junction equivalent. The values in the BigWig files are stored as the number of reads overlapping each base pair so for each feature, we take the sum of these values and then divide by the read-length of the experiment to determine the equivalent number of reads overlapping each feature. No such normalization is necessary for the values from the junction coverage file.

### Deriving model inputs

Our goal is to derive transcript abundances given the known gene structure and feature counts summarized above. Continuing with the example gene from **Figure 1**, transcript 1 is composed of all 7 features, while transcript 2 is composed of features 1 and 2 only. Based on this structure, reads aligning to features 3-7 should have originated from transcript 1 and not transcript 2. We use this structure to set up the design matrix for our linear model.

We name our independent variables **feature probability** vectors, and denote them as *X*^1^ and *X*^2^ respectively for each transcript. Each vector is of length 7, corresponding to the number of features. Each element 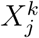 encodes the probability a random read from transcript *k* overlaps feature *j* of our gene, where *k* = 1 *…* 2 and *j* = 1 *…* 7. The column-wise collection of feature probability vectors for our example gene is denoted as **X** with dimension [7×2] and is referred to as the **feature probability matrix** for this gene. Note that the values in this matrix depend on both the calculated features and the length of the reads of the sequencing experiment (100bp in this example).

The true **X** is not known, but it can be estimated based on sequence content of the transcripts and the read length of the experiment. To estimate *X*^1^, sliding segments of 100bp from transcript 1 are aligned to the GRCh38 reference using the aligner HISAT2 [13]. The number of aligned segments overlapping each feature is summed and divided by the number of total 100bp segments to produce the estimate of *X*^1^, denoted by 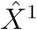. The estimated feature probability matrix 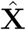 is the column-wise collection of such estimated feature probability vectors for all transcripts in the gene. More complex implementations can readily include adjustments for GC content, 5’ bias, and mappability differences by weighting each row of 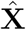 appropriately.

### Non-negative linear model

For our gene, we denote the observed feature counts vector *Y*. The underlying assumed data generation process is illustrated in **Figure 7**. We are interested in estimating the true transcript abundances *β* using our estimated 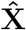 by solving for:

**Figure 7:**
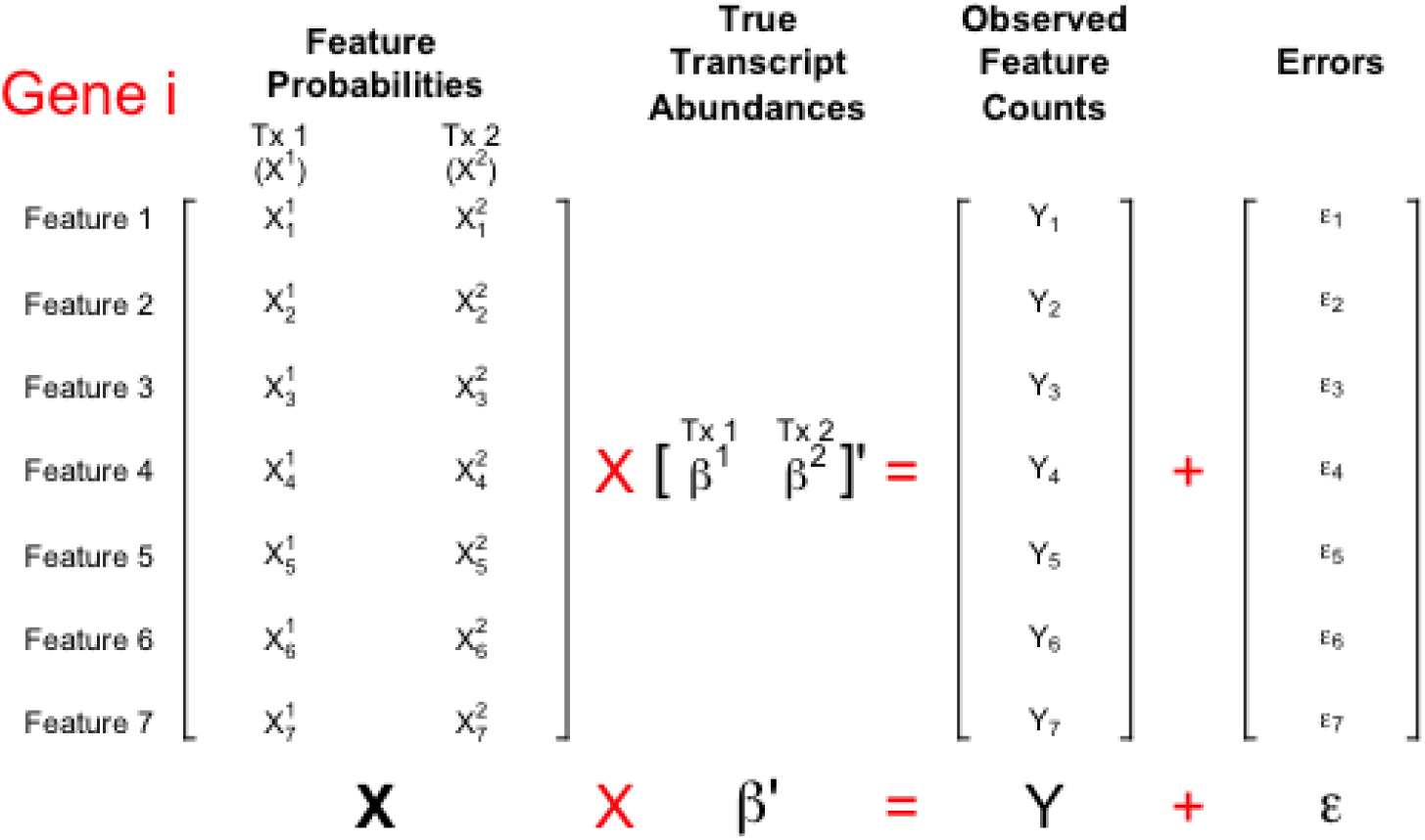
For one example gene, an illustration of our model formulation of the relationship between transcript abundances and observed feature counts. A column of the feature probability matrix represents the expected contribution to the observed feature counts by a random read from the corresponding transcript. Our model estimates *β* using a non-negative linear model. Furthermore, since the true feature probability matrix is unknown, we estimate it by applying the aligner HISAT2 to possible reads from each transcript.

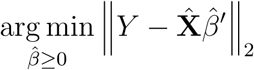

subject to the constraint that each element of 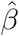 is non-negative. Many existing algorithms and implementations exist for finding the solution. For recountNNLS, we used the function nnls found in the R package nnls [19] for ease of implementation.

### Standard error calculation

Our model is amenable to standard error estimation of 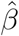 using a heteroscadastic consistent sandwich estimator proposed and referred to as HC4 by Cribari-Neto [20]. The covariance matrix of 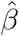 is estimated using our feature probability matrix **X** as:

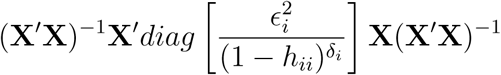

where *∈* _*i*_ is the residual from the *i*-th feature, and *h*_*ii*_ is the *i*-th diagonal of the projection (“hat”) matrix **H** calculated as:

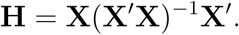

We define *δ*_*i*_ = *min*(4, *n * h*_*ii*_*/p*), with *n* the number of transcripts and *p* the number of features at a given locus. *h*_*ii*_ is capped to be at most 0.99 to ensure division by 0 does not occur for points with *h*_*ii*_ computationally equal to 1.

For transcripts that only have 1 feature while being quantified, the reported standard error is 0 and reflects that one is unable to quantify the variability of the estimate.

### Confidence interval construction

The construction of confidence intervals is based on a t-statistic approach, where the critical value depends on a degree of freedom equal to *n - p*. *n* is the number of features and *p* is the number of transcripts being estimated at once. If we let 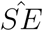 denote the diagonal of the estimated covariance matrix for 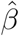, the *α*-level confidence intervals are:

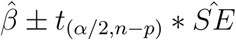

### Uniqueness score

Our method also produces a uniqueness score for transcript *i*, defined as *rss*_*i*_*/tss*_*i*_. *rss*_*i*_ is the residual sum of squares for transcript *i* after fitting *NNLS*(**X**_*i*_, **X**_*-i*_), where **X**_*i*_ is the *i*th column of our feature probability matrix **X** and **X**_*-i*_ is **X** with the *i*th column removed. *tss*_*i*_ is the sum of the squares of each term of **X**_*i*_.

### Quantification compilation for recount2

To quantify the entire transcriptome for a given sample, we execute separate linear models on each of what we refer to as “bundles” of genes. We define a single bundle as all genes that share any non-zero entries in the feature probabilities of transcripts associated with those genes. As different read lengths affect the multi-mapping of sequencing reads, we calculate the set of bundles for each read length, by examining all feature probability matrices. However, for read lengths of 37bp and 50bp, this results in the creation of a bundle encompassing 1966 and 4953 genes respectively too large to handle in computation at once. As such, for 37bp and 50bp datasets, we resort to approximating the bundles with the bundles built for 75bp, resulting in some increase in bias, but allowing for computational tractability. As an addition to the recountNNLS package, we offer precomputed features, feature probability matrices, and bundles reflecting read lengths of 37, 50, 75, 100, and 150bp in the package recountNNLSdata.

Our method produces a Ranged Summarized Experiment (RSE) object per project mirroring the structure of recount2. For each sample of a project, we utilize the set of feature probability matrices that match the read length of the sample most closely. If the match is not exact, we adjust the estimated abundances and standard errors by the ratio of feature probability matrix read length over actual sample read length.

For each project, the RSE object contains the estimated fragment counts, standard errors, uniqueness scores, and degrees of freedom, accessible via the function assays as fragments, ses, scores, and df respectively. Each row of the fragments, ses, scores, and df matrices represents a transcript, and each column represents a sample. The corresponding transcripts are stored as a GRangesList accessible via the rowRanges function, and meta information such as length and number of exons is stored in a table accessible by the function rowData. Transcripts that introduce colinearity (either perfect or computational) in the model matrix 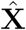 are reported as NA in counts and ses. Transcripts are deemed too colinear by default behavior of the *lm*() function in *R*.

### Performance evaluation

Using our linear model on real and simulated data, we compare our estimates to those from the established methods Kallisto [7], Cufflinks [13], RSEM [9], and Salmon [8].

#### Dirichlet-negative binomial simulation scenarios

We simulated RNA-seq data using the R package polyester [14] under 10 scenarios: read lengths of 37, 50, 75, 100 and 150bp with either single-end or paired-end FASTA reads. For the sake of simulation expediency, we selected all coding transcripts from chr1 and chr14 from the GencodeV25 transcriptome annotation, which comprises 12.5% of the entire annotation (18,303 total transcripts). The reads are generated via polyester [14] with a Gaussian fragment length distribution with mean 250 and standard deviation 25. The number of reads to simulate was determined on a gene-by-gene basis, with most genes having a dominant transcript that produces over 50% of the sequencing reads. The relative abundances of the transcripts are chosen via a Dirichlet distribution with *α* = 1*/f*, where *f* is the number of transcripts in the gene. The total number of reads at a gene is chosen as a negative binomial with size=4 and p=0.01. The number of reads of each transcript is the product of the outcomes of the Dirichlet and the negative binomial. These parameters produce on average 400 fragments for each of the 2,675 genes, distributed to approximately 60 fragments per transcript.

We created alignment indices for the subset of the transcriptome from chr1 and chr 14 for use with Kallisto [7] and Salmon [8]. The simulated FASTA files were fed to Kallisto [7], HISAT2-Cufflinks [13, 6], RSEM [9], and Salmon [8] with default parameters where applicable. Methods were only asked to quantify the abundances of the subset of transcripts from protein-coding genes on chr1 and chr14. For single end simulations, Salmon [8] and Kallisto [7] require input of the fragment length distribution, for which the true parameters of (250, 25) were used. For Cufflinks [6], we provided the fragment length ditsribution, and used --total-hits-norm --no-effective-length-correction --no-length-correction options. For our linear model, we utilized Rail-RNA [5] to process the FASTA files in the same manner as in recount2 [4]. For evaluation, each method’s abundance estimates (*est*) were compared to the true number (*truth*) using mean absolute error (MAE):

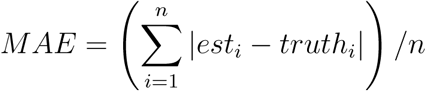

where *i* denotes a transcript out of the *n* = 18, 303 transcripts selected from chr1 and chr14. For each scenario, approximately 51% of the 18,303 transcripts had true non-zero expression.

#### Confidence interval coverage by transcript

We randomly sampled 2000 transcripts from the set of transcripts belonging to protein-coding genes from chr1. We simulated 100 repeated datasets for each Dirichlet-negative binomial scenario described above, with each selected transcript receiving 20x coverage. The simulated fasta files were aligned via Rail-RNA, and the output BigWig and junction files were passed to our model for quantification. Confidence intervals were constructed using the t-statistic-based method described above. For a given transcript, the number of simulations in which the confidence interval for that transcript covered the truth was recorded.

#### Hybrid simulation scenario

Using polyester, we also simulated a dataset with 75bp read length and paired-end reads to mimic the expression levels observed in sample ERR188410 of the GEUVADIS Consortium dataset. The ground truth number of counts generated for each transcript was taken from the estimated counts from applying RSEM to transcripts that are a part of protein-coding genes in the GencodeV25 annotation. Approximately 50% of the 144,729 transcripts reported by RSEM received a positive count. Although these counts may not be the truth, they might better capture patterns of correlation and variability present in real data than the simulation scenario described above did. The insert length was again set to have a mean of 250 and a standard deviation of 25. The simulated FASTA files were used as input for Salmon [8], Kallisto [7], and HISAT2-Cufflinks [13, 6], while the Rail-RNA output BigWig and junction files were used as input for out method. We asked each method to quantify all transcripts from protein-coding genes of the GencodeV25 annotation, using suitable indices for each method built on the entire GencodeV25 annotation. Cufflinks [6] was again used with --total-hits-norm --no-effective-length-correction --no-length-correction options. For evaluation, each method’s estimates were again measured using MAE for the full set of transcripts belonging to protein-coding genes across all chromosomes.

### GEUVADIS Consortium

We downloaded the raw paired-end FASTQ files for sample ERR188410 of the GEUVADIS Consortium dataset. The FASTQ files were used directly as input for Kallisto [0.43.0] [7], HISAT2 [2.0.5]-Cufflinks [2.2.1] [13, 6], RSEM [1.3.0, bowtie 1.1.1] [9], and Salmon [0.8.2] [8] using default parameters. The recount2 summary measures for the GEUVADIS project samples were used as inputs for our linear model. We were only interested in estimating the abundances of the transcripts belonging to protein-coding genes in the GencodeV25 annotation. Indices were built for the GencodeV25 transcriptome where needed. Cufflinks [6] was run with --total-hits-norm --no-effective-length-correction --no-length-correction. Abundance estimations on the transcript- and gene-level were compared pair-wise between methods using Spearman’s correlation. We also examined pair-wise the number of transcripts that were assigned non-zero expression by both methods.

## Declarations

### Ethics approval and consent to participate

Not applicable.

### Consent for publication

Not applicable.

### Availability of data and material

The following code will estimate transcript level abundances and associated standard errors for a given project in the recount2 repository) (if R has access to sufficient resources) for a given project id. An example case is demonstrated below for project DRP000366, and additional commands are located in the supplement.

~~~
devtools::install_github(’JMF47/recountNNLSdata’, ref=’4202515’)
devtools::install_github(’JMF47/recountNNLS’, ref=’9acad0b’)
library(recountNNLS)
pheno = processPheno(’DRP000366’)
rse_tx = recountNNLS(pheno)
~~~

The scripts to reproduce the analyses and plots in this manuscript are available at https://github.com/JMF47/recountNNLSpaper

### Authors’ contributions

JTL, JMF, KK, and MAT envisioned the objectives of the study. JMF derived the model with input from all other authors. JMF implemented the R package with contribution from LCT. JMF, JTL, KK, and MAT wrote the manuscript. All authors read and approved the manuscript.

## Acknowledgements

We would like to thank SciServer for hosting the recount2 files. SciServer is being developed at, and administered by, the Institute for Data Intensive Engineering and Science at Johns Hopkins University and is funded by the National Science Foundation Award ACI-1261715. For more information about SciServer, visit http://www.sciserver.org/.

